# Neural computations underpinning the strategic management of influence in advice giving

**DOI:** 10.1101/121947

**Authors:** Uri Hertz, Stefano Palminteri, Silvia Brunetti, Cecilie Olesen, Chris D Frith, Bahador Bahrami

## Abstract

Research on social influence has mainly focused on the target of influence (e.g., consumer, voter), while the cognitive and neurobiological underpinnings of the source of the influence (e.g., spin doctor, financial adviser) remain unexplored. Here, we introduce a 3-sided Advising Game consisting of a client and two advisers. Advisers managed their influence over the client strategically by modulating the confidence of their advice depending on their level of influence (i.e. which adviser had the client’s attention) and their relative merit (i.e. which adviser was more accurate). Functional magnetic resonance imaging showed that these sources of social information were computed in distinct cortical regions: relative merit prediction error was tracked in the medial-prefrontal cortex and selection by client in the right temporo-parietal junction. Trial-by-trial changes in both sources of social information modulated the activity in the ventral striatum. These results open a fresh avenue for exploration of human interactions and provide new insights on the neurobiology involved when we try to influence others.

## Introduction

It is hard to overstate the role of social influence in our lives. From the outcomes of political campaigns to our stand on issues such as global warming, immigration and taxation, the fate of such critical matters in our public lives depends on persuaders working to influence public opinion^1^. Social influence is also intertwined with our everyday life, as we try to persuade our children, influence our bosses, and gain popularity among our friends. Research on social influence has been dominated by the motivation to understand the minds of the targets of influence (e.g., consumers, voters) in order to exert even more influence on them^2^. Far less is known about the cognitive and neurobiological processes at play in the persuaders (e.g. spin doctors, financial advisers). Here we ask what happens in the persuader’s mind when engaged in the attempt to influence others.

Bayarri and DeGroot^3^ proposed a normative solution for how advisers should modulate their strategy for offering their predictions in order to maximise their influence. They assumed that clients are affected by advisers’ accuracy and confidence^4–7^, i.e. they will be more likely to follow advisers that express confidence only when it is warranted, and discredit highly confident but inaccurate advisers. In this case, in order to be selected by the client, advisers should modulate their advice giving strategy depending on their current influence on the client. (a) When the adviser is failing to influence the advisor, s/he should express higher confidence than permitted by objective evidence (positive advice confidence deviance). (b) When the adviser is trying to maintain and protect already high influence, s/he should sit on the fence with cautious, nuanced, advice that is lower than justified by the evidence (negative advice confidence deviance). We will call this a ‘competitive’ strategy for acquiring and maintaining influence.

Social rank theory^8,9^ proposes an alternative account of advising behaviour by positing that people are not motivated just by the desire to influence others’ choices, but also by the fear of being excluded by their target. This theory suggests that an adviser’s confidence should be proportional to her rank in a group. Lower rank individuals adopt submissive behaviours, expressing lower confidence and avoiding eye contact^8^. Accordingly, social rank theory suggests that, in contrast to the ‘competitive’ strategy described above, humans may adopt a ‘defensive’ strategy to manage influence by giving cautious advice when they are ignored by the client (i.e. when their influence is low) and exaggerating their confidence when their influence is high. In addition, social rank theory underscores the importance of an active process of social comparison by which an adviser evaluates her rank by tracking her performance relative to rival advisers ^10–14^. In this view, relative performance or merit may also affect advice confidence leading people to display higher confidence when they think they perform better than their peers.

These theoretical models of strategic advice giving and influence management may rely on a number of social cognitive processes, e.g. mentalizing (theory of mind)^15^, social motivation^16^, and social comparisons^17^, which have been previously linked to specific neural substrates^16,18^. To track one’s current level of influence (i.e. client preference for each advisor) the adviser needs to evaluate the client’s state of mind from his actions, presumably incorporating the brain’s “theory of mind” (ToM) ^17^ apparatus. ToM processes, such as mentalizing and updating other’s beliefs, have been linked to activity in the right Temporo-Parietal Junction (rTPJ)^19,20^ and the medial Prefrontal Cortex (mPFC)^15,20–23^. Influence management also requires the adviser to track her own and her rival’s performance in order to calculate her relative merit and adjust this as new evidence about rival’s performance emerges ^13^. Previous works suggest that this cognitive process is linked to activity in the mPFC, which tracks rank and social status ^24,25^ and the performance of collaborators and competitors ^26–29^. Finally, the adviser is motivated to increase her influence over the client and her merit over the rival adviser ^13^. Brain areas such as the Ventral Striatum (VS) and the ventro-medial Prefrontal Cortex (vmPFC) ^30^, that track primary (e.g. food) rewards, are also responsive to secondary social rewards such as being selected by others ^31^, having an increase in social status ^25^ and being confirmed by the group ^32^. These studies lead us to hypothesise that fluctuations in social status and relative merit are tracked by a reward-sensitive network.

Here we examine how people strategically manage their influence, and test the hypotheses we derived above regarding the cognitive processes and neural systems underlying this process. First, we examined the behavioural predictions of a normative model of advice giving ^3^ against those drawn from social rank theory ^8^. We asked if people would give overconfident (‘competitive’ strategy ^3^) or under-confident (‘defensive’ strategy ^8,9^) advice when they are ignored by their client, and whether social comparison with a rival adviser plays a role in advising behaviour. We then sought to identify the neural mechanisms underlying the participants’ attempts to influence others by advice giving, examining whether (and how) brain areas associated with mentalizing, social comparison and valuation track the appraisal made by a client and calculate relative performance during a strategic advice giving task.

## Results

### Behavioural Task

We devised a social influence scenario in which two advisers competed for influence over a client (Figure 1, online demo: http://www.urihertz.net/AdviserDemo). On a series of trials, a client is looking for a reward hidden in a black or a white urn. He relies on two advisers who have access to evidence about the probability of the reward being in the black or the white urn. At the beginning of each trial (appraisal stage), the client chooses the adviser whose advice (given later) will determine which urn the client will open. The client’s choice of adviser is displayed to the advisers, who then proceed to the evidence stage. They see a grid of black and white squares for half a second. The ratio between the black and white squares indicates the probability of the reward location. Next step is the advice stage. Each adviser declares her advice about the reward location using a 10-level confidence scale, ranging from ‘certainly in the black urn’ (5B) to ‘certainly in the white urn’ (5W). Subsequently, both declarations are shown to both advisers and the client (showdown stage). Finally, at the outcome stage, the urn indicated by the chosen adviser’s advice is opened, and its content is revealed to everyone. The next trial begins with the client selecting an adviser – the client can decide to switch advisers or to stick with the same adviser.

**Fig. 1:**
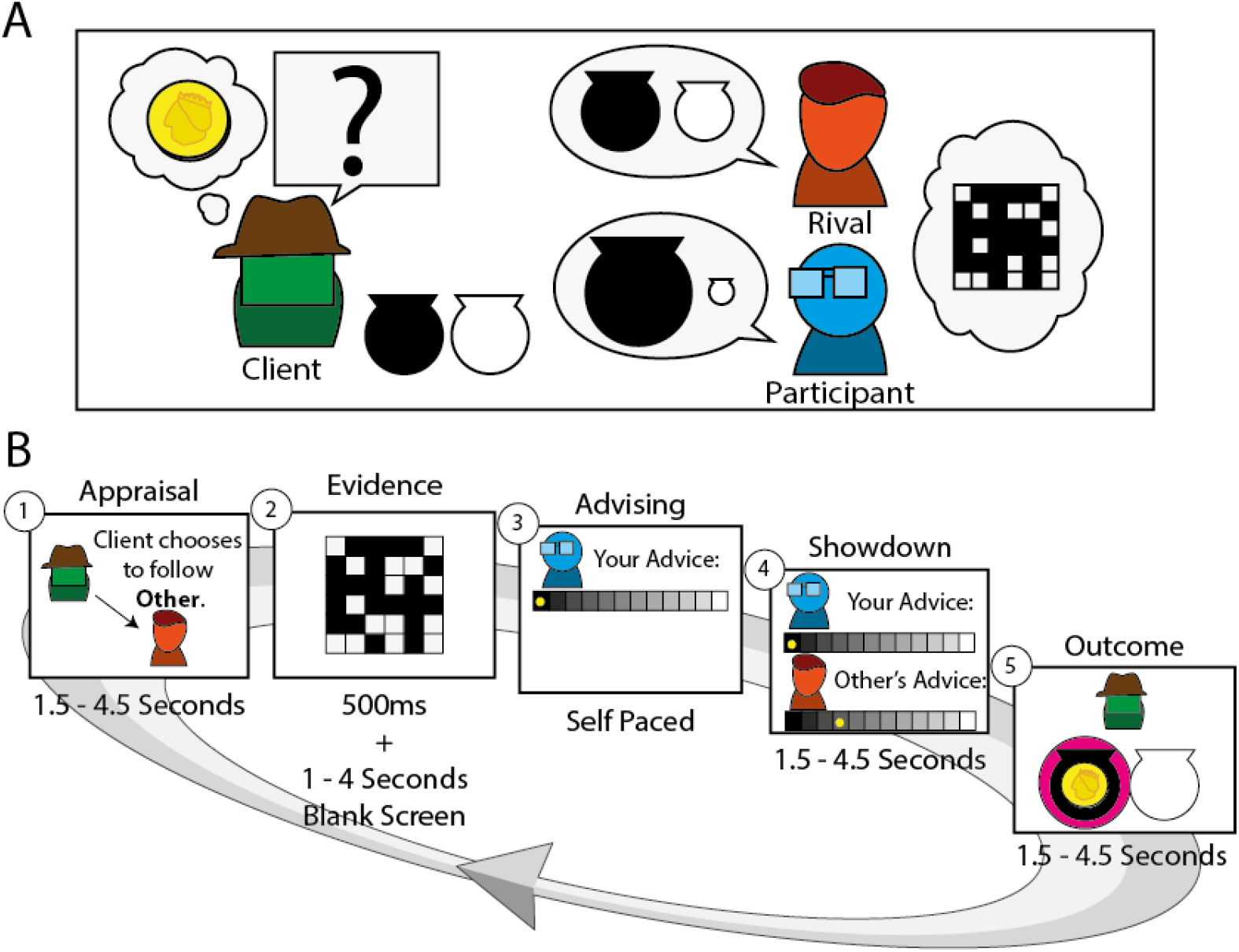
Experimental Design. (A) Participants were engaged in an advice giving task. In this task a client is looking for a coin hidden in a black or white urn. He relies on advice from two advisers (the participant (blue) and a computer generated adviser (red)). The advisers, but not the client, have access to information regarding the probability of the coin location. The client considers the advisers’ previous success and current confidence when choosing an adviser to follow on each trial. (B) Each trial contains five stages. (1) Appraisal: In the beginning of each trial the client chooses the adviser he is going to follow on the commencing trial (and consequently which adviser is ignored). (2) Evidence: The participant (and the rival) then sees a grid of black and white squares, whose ratio represents the probability of the coin being in the black urn. (3) Advising: The participant states his advice on coin location using a 10 levels confidence scale ranging from “definitely in the black urn” (5B) to “definitely in the white urn” (5W). (4) Showdown: both advisers’ opinion is displayed. (5) Outcome: The content of the urn suggested by the selected adviser (magenta circle) is revealed. The next trial starts with appraisal by client based on the history of confidence and success. The stages timings indicated are from the fMRI experiment, where jitter was introduced between stages. See supplementary materials for online and lab stages timings. An interactive demo of the experiment can be viewed at http://www.urihertz.net/AdviserExperiment.html.

We were interested in the way advisers use advice confidence as a persuasive signal to manage their influence over the client. In our first experiment, therefore, all participants were assigned to play the role of *adviser*, while the rival adviser and the client’s behaviour were governed by algorithms adapted from Bayarri and DeGroot^3^ (see Methods). We examined participants in three independent cohorts: online (N=58), in the lab (N=29) and in-the-scanner (N=32). There were some minor differences between the groups, as online participants performed fewer trials than those in the lab and the scanner. Online participants were paid a bonus for the number of times the client selected them, while the cohorts tested in the lab and the scanner received a fixed monetary compensation (see full details in Methods section). We observed no difference in the 3 cohorts’ behaviour. Finally, to examine whether the observed behaviour can be generalised to real life social interactions, we ran another lab-based interactive version of the experiment (48 participants organized in 16 triads, thus comprising 32 advisers and 16 clients) in which all roles (advisers and client) were played by participants who genuinely interacted with one another.

### The effect of Selection by Client and Relative Merit on advising

To assess the advising behaviour, we examined the trial-by-trial deviance of advice confidence from probabilistic evidence (Figure 2A). If the adviser is strictly committed to communicating the information she is given, then confidence exactly matches the ratio of black to white squares in the evidence grid (Figure 1B). For such adviser confidence level ‘5B’ is reported when close to 100% squares in the grid are black indicating that probability of the coin being in the black urn is almost certain. Conversely, confidence ‘1B’ stems from weak evidence and indicates the probability of the coin being in the black urn is around chance (50%). Advice confidence would deviate if the confidence is higher (positive deviance) or lower (negative deviance) than the probability indicated by the evidence. Participants’ advice deviance was significantly greater than 0 (t(119) = 17.08, p < 0.00001), as they displayed systematic overconfidence in their advice. Importantly, participants’ advice deviance was affected by the ‘Selection by Client’ parameter (ignored vs chosen), and was larger, i.e. more overconfident, when ignored by the client compared to trials in which the participant was the selected adviser (paired t-test, t(119) = 3.1, p = 0.002) (Figure 2B). Moreover, participants adjusted their advising policy dynamically: overconfidence increased in periods when they were being ignored (not chosen by client) and was attenuated during periods of being selected by the client (Figure 2C).

**Fig. 2:**
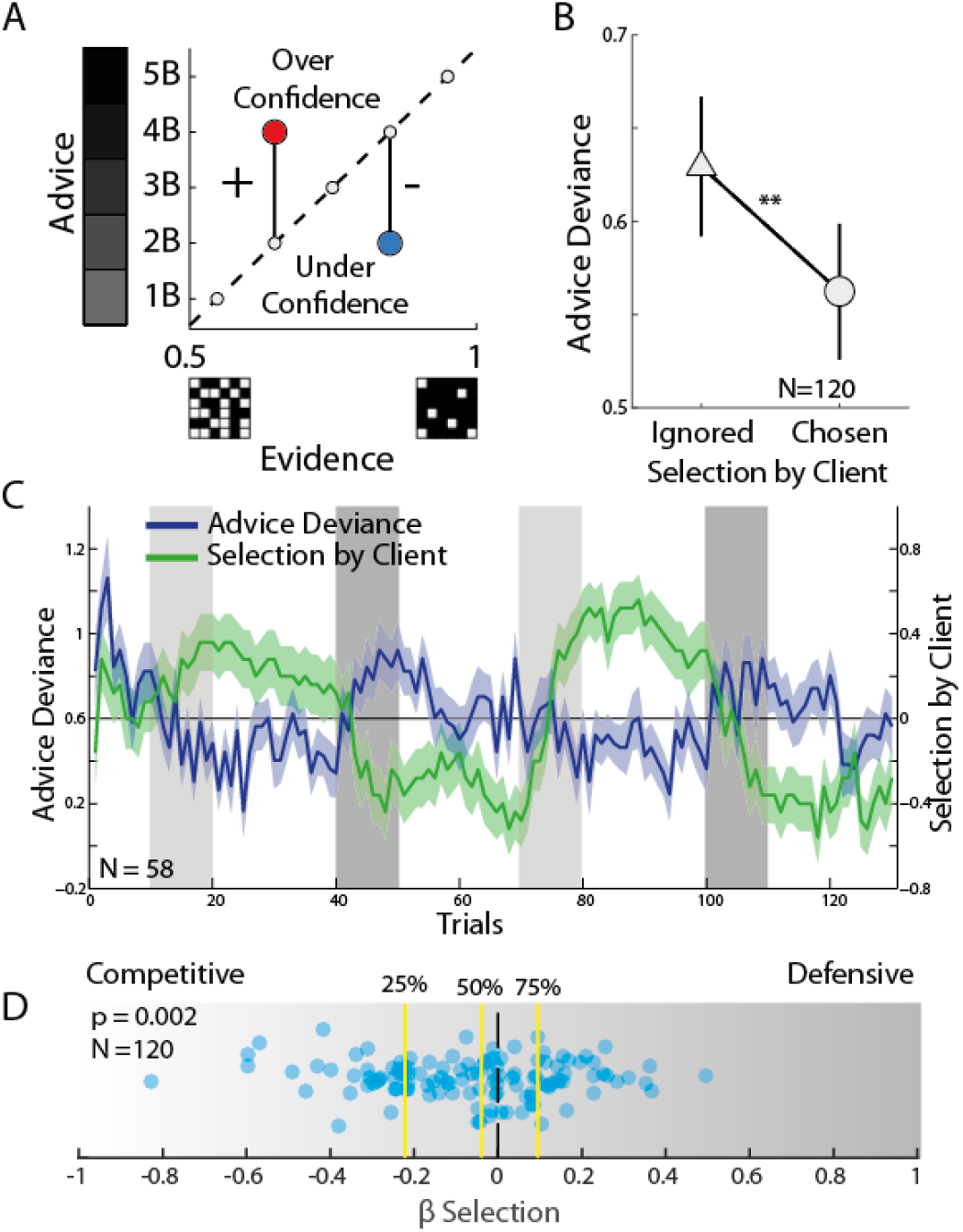
Selection by Client Effect on Advice. (A) Deviance of advice from probabilistic evidence. Confidence is plotted against evidence grid, i.e. probability that coin is in the black urn. The dashed line represents zero-deviance policy in which confidence matches the ratio of black to white squares in the evidence grid. Overconfident advice (red circle) would lie above the dashed line and was defined as positive deviance; conversely, underconfident advice (blue circle) corresponded to negative deviance. (B) Averaged advice deviance under high vs low selection (paired t-test, t(119) = 3.10, p = 0.002). (C) The dynamic interplay of selection and advice deviance in the online experiment. Blue line = trial-by-trial average advice deviance. Green line = average selection. Light green and blue = standard error of the means. Light and dark grey blocks mark periods of low evidence reliability for the rival and for the participant, respectively (see Fig. S1). (D) Our best fitting model contained a parameter governing how Selection by Client affect advice deviance, ‘β-Selection’. Positive values of β-Selection are associated with the defensive strategy, i.e. increased advice deviance when being selected. Negative values are associated with the competitive strategy, i.e. decreased advice deviance when being chosen by the client. Despite high level of individual differences, this parameter was significantly lower than 0 across participants (Mean±SEM = -0.08±0.02, t(119) = 3.12, p < 0.002), in line with our previous analyses.

This finding demonstrated how being selected over a rival shapes the attempt to influence the client, in line with the predictions of the competitive strategy. However, our analysis so far focused only on selection by the client, i.e. the current level of influence on the client, as the force shaping our attempt to influence others, while ignoring the effect of social comparison with the rival adviser implicated in social rank theory ^10–14^. A deeper understanding of the participant’s advising strategy would require factoring in the interaction between the (exogenous) Selection by Client and the (endogenous) Relative Merit. To disentangle the impact of Selection by Client and Relative Merit on behaviour we employed a computational approach.

While trial-by-trial Selection by Client was made public to the participants and explicitly available for analysis, trial-by-trial variations of Relative Merit had to be inferred to examine its effect on advice giving. To this end we followed the model fitting approach used in behavioural and neuroimaging studies to estimate the latent subjective processes that involve adviser’s reliability, reward prediction errors and social comparisons ^29,33,34^. We devised several models that made different assumptions about how Relative Merit is computed and how it may affect advice deviance and formally compared the models using the empirical data. The winning model’s estimates of trial-by trial Relative Merit were used to interpret the advising behaviour. We noted that participants could evaluate the prognostic value of their own and their rival’s advice every time the outcome (i.e. the content of the chosen urn) was revealed (Figure 3A). For example, strong advice for the black urn (i.e. 5B) would have better prognostic value than a cautious advice supporting the same choice (i.e. 1B) if the black urn turned out to produce a coin. Conversely, advising 1W would have better prognostic value than 5W if the white urn was chosen but turned out to be empty. Prognostic value is calculated by multiplying advice confidence with its accuracy (correct = 1, wrong=-1) (i.e. whether the urn suggested by the advice contained a coin, see methods for further details). On a given trial *t*, we update the Relative Merit variable by using the difference between the prognostic value of the participant’s and the rival’s advice such that:

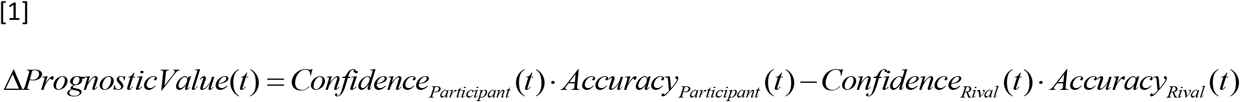

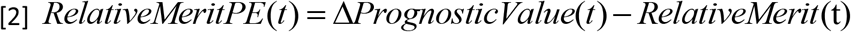

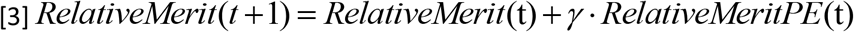

In equation [3] *γ* is the learning rate of change to Relative Merit. Relative Merit is positive if the participant’s advice had consistently higher prognostic value compared to rival’s advice. Importantly, Relative Merit is calculated independently from selection by the client and gives a quantitative estimate of the latent subjective process of social comparison.

To quantify whether and how this measure of Relative Merit could explain overconfidence behaviour, we fitted a set of hierarchically nested computational models to advice deviance. Our most simple model had only one free parameter for systematic bias in advising, capturing the average (intercept) advice deviance, which stands for trait overconfidence and under-confidence (Bias Model). Our next model included the trial-by-trial selection by the client as well (Client Model). Positive values of the weight assigned to Selection by Client weight, would indicate that the participant followed a ‘defensive’ strategy, expressing higher confidence when selected by the client. Conversely, negative values for this parameter would correspond to the ‘competitive’ strategy. These models should account for the effects observed in our initial analysis, and therefore can serve as a baseline to compare more elaborate models that also include a Relative Merit measure. Our next model included the sign of the Relative Merit from equation [3] (Mixed model). Finally we set up a model which included all of the above and the interaction between Selection by Client and Relative Merit (Interaction model, Equation 4]). We also fitted models that used the sign and amplitude of Relative Merit (see Supplementary Materials).

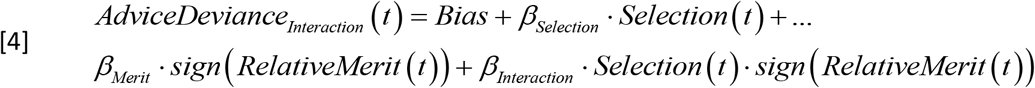

After fitting all models to the advice deviance data and compensating for the number of free parameters (see Supplementary table ST1, Supplementary Figure S2 and S3), we found that the interaction model (Equation [3]) gave the best fit to the empirical data. Our model fitting procedure estimated individual parameters for *Bias, γ, β_Selection_*, *β_Merit_* and *β_Interaction_* (Supplementary Figure S3, Supplementary table ST1). Bias parameter was significantly higher than zero across participants, verifying our observation that participants were generally overconfident (Mean ± SME Bias: 0.67 ± 0.05, t(119) = 13.1, p < 10^−10^). In addition, the selection parameter (*β*_*Selection*_) was significantly lower than zero across participants (Mean ± SME: -0.07± 0.02, t(119) = -3.2, p = 0.002), supporting our direct comparison in Figure 2B, favouring the ‘competitive’ over ‘defensive’ hypothesis.

While selection parameter (*β_Selection_*) was significantly lower than 0, we observed high degrees of variability among individuals, with a positive selection parameter estimated for one third of our participants (38) (Figure 2D). Previous works on social rank theory ^8,35^ suggested that negative self esteem, i.e. seeing oneself as inferior to others and less desirable, may lead to displays of low confidence in social interactions and a greater receptivity to negative social signals. We reasoned that population variability in behavioural response to social selection and/or exclusion (as quantified here by our selection parameter) may be accounted for by individual differences in receptivity to negative social feedback. To test this hypothesis, we went back to our pool of participants and were successful at recruiting N=69 of them to complete the Fear of Negative Evaluation questionnaire (FNE) ^8,36^. We found that FNE score correlated with the participants’ model estimate of selection parameter, as participants with higher FNE score, i.e., more negative self-perception, were more likely to follow the defensive strategy (N=69, R = 0.25, R^2^ = 0.06 p = 0.036). By linking previous research on social rank and our participants’ behaviour, this latter finding provided evidence of external validity for our computational model.

Estimated model parameters associated with Relative Merit, and the interaction term between Relative Merit and Selection by Client were harder to interpret directly. Even greater individual differences were observed for these parameters. When averaged across participants, neither parameter was significantly different from zero (t(119) = < 0.7, P > 0.45), with individuals varying in the effect of Relative Merit and Interaction (see Table ST1). However, model selection and comparison showed beyond doubt that the interaction parameter did provide the model with better power to capture behaviour. To examine the contribution of all parameters to the explanation of the behavioural data, we used the sign of the Relative Merit estimated by the interaction model using the fitted Relative Merit learning rate *γ* (Equation [3]), to label the trials as ‘positive’ vs ‘negative’ Relative Merit. Each trial was also categorised according to the Selection by the Client as ‘ignored’ or ‘chosen’. We then examined advice deviance across the resulting 2x2 combinations of Relative Merit (positive vs. negative) and Selection by Client (selected vs. ignored) using a repeated measures ANOVA. We found a significant effect of Relative Merit (F(1,358) = 14.4 p = 0.0002), a significant effect of Selection by Client (F(1,358) = 8.1, p = 0.005) and a significant interaction effect (F(1,358) = 17.95, p < 0.0001) on advice deviance, as model fitting procedure had suggested (Figure 3B, see Figure S5 for breaking of these results across cohorts). Importantly, this analysis elucidated the nature of the interaction by showing that participants expressed significantly greater confidence in their advice on trials in which they assumed they had done better than their rival (i.e. positive Relative Merit) but nonetheless had been (perhaps unexpectedly) ignored by the client. To put it metaphorically, advisers shouted most loudly when they had reason to believe that their merits had been overlooked. We used a simulation to examine whether such pattern could be recovered by one of our competitive models^37^, and found that only the interaction model (equation [4]) could reproduce this pattern of results (Figure S4).

**Fig. 3.**
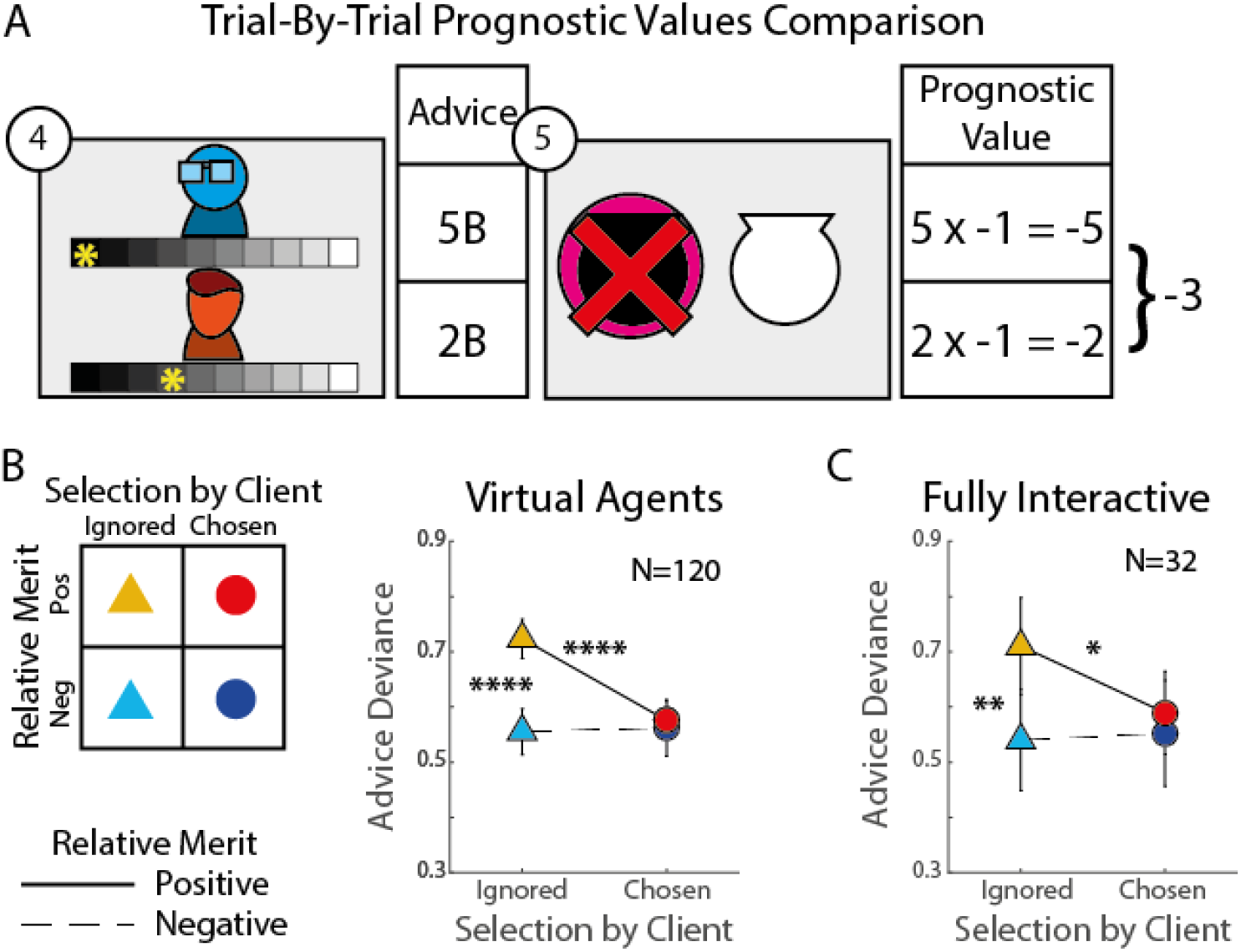
Relative Merit Effect on Advice. (A) At the end of each trial participants could evaluate and compare their and the rival’s advice prognostic value. The difference in prognostic value was accumulated to form the participants’ Relative Merit. (B) Our best fitting model included a free parameter for accumulation rate for differences in prognostic values, for Relative Merit and for the interaction between merit and client’s selection. We used the individual estimated parameters to evaluate the trial by trial Relative Merit, and divided the trials to four conditions according to the Selection by Client and Relative Merit variables. Advice deviance was highest (i.e. most overconfident) when participants’ positive Relative Merit conflicted with the client’s choice to ignore them, and demonstrated a significant interaction effect. (C) The same pattern of behaviour was replicated in the fully interactive experiment, where three human participants played the roles of a client and two advisers. (two-tailed paired t-test comparisons: * p < 0.05, ** p < 0.005, **** p < 0.00005, Error bars indicate SEM).

The behaviour we observed (Figure 3), one may argue, can arise in response to the specific manner in which our algorithms, following Bayarri and DeGroot’s assumptions^3^, controlled the client and rival adviser behaviour. To confirm the validity and generality of our results, we ran a fully interactive experiment in which both the advisers and the client were human participants (see Methods for details) and no agent’s behaviour was under experimenter control. The scenario was the same as before, but now involved three participants engaged in a multiplayer game played on three computers in three adjacent cubicles connected via the internet. We applied the same analysis and model fitting and estimation of Relative Merit and Selection by Client effects to the advisers’ behaviour in this fully interactive experiment. The results replicated the main results from the virtual agents’ experiment: advice deviance was highest when the adviser was ignored but her Relative Merit was positive (Figure 3C; mixed effects ANOVA; a significant interaction effect (F(1,96) = 5.05, p = 0.03, and a significant effect of Relative Merit, F(1,96) = 4.43, p = 0.04). In addition, we did not observe any significant discrepancies between live client’s behaviour and virtual client behaviour (Figure S6).

To further examine the relation between the virtual and live experiments, we fitted data from all experimental cohorts, within a mixed-effect ANOVA with Relative Merit (positive/negative), Selection by Client (chosen/ignored) and Agents (live/virtual) as main factors and subjects as random effect factor nested within the Agents factor. We did not observe any effect of Agents on advice, neither in the main effects nor the interaction. Overall, we observed a significant main effect of Relative Merit (F(1,442) = 13.73 p = 0.0004), a significant main effect of Selection by Client (F(1, 442) = 5.61, p = 0.019) and a significant interaction between the two factors (F(1, 442) = 14.16, p = 0.0002).

The results from the fully interactive experiment provided an additional replication of our main results, demonstrating the robustness of the findings and strengthening the ecological validity of the paradigm in generalizing the findings to interactive human behaviour.

### Neural Correlates of Selection by Client and Relative Merit

Having established the effects of Selection by Client and Relative Merit on advising behaviour, we used the same paradigm to examine the neural correlates of the attempt to influence others. Employing functional magnetic resonance imaging (fMRI), each of the five stages of a trial was treated as an event in a fast event-related design (see Methods). Using the computational model-based analysis described above, we labelled trials according to their Relative Merit (positive vs. negative), and Selection by Client (ignored vs. chosen). The resulting 2x2 combination of conditions allowed us to examine the changes in the blood oxygenation level dependent (BOLD) signal that correlated with the main effects of selection and its interaction with Relative Merit. Activity in the right Temporo-Parietal junction (rTPJ) (p < 0.001, FWE cluster size corrected p < 0.05) (Figure 4 and Table ST2) displayed the main effect of Selection by Client (Ignored > Selected): BOLD signal was higher when the participant was ignored (vs. selected) by the client (regardless of the sign of Relative Merit) during the observation of evidence. This finding is in line with previous studies showing increased activity in the rTPJ when others’ actions did not match the participant’s predictions ^38^, and when inferring others’ intentions from their actions ^20^.

**Fig. 4:**
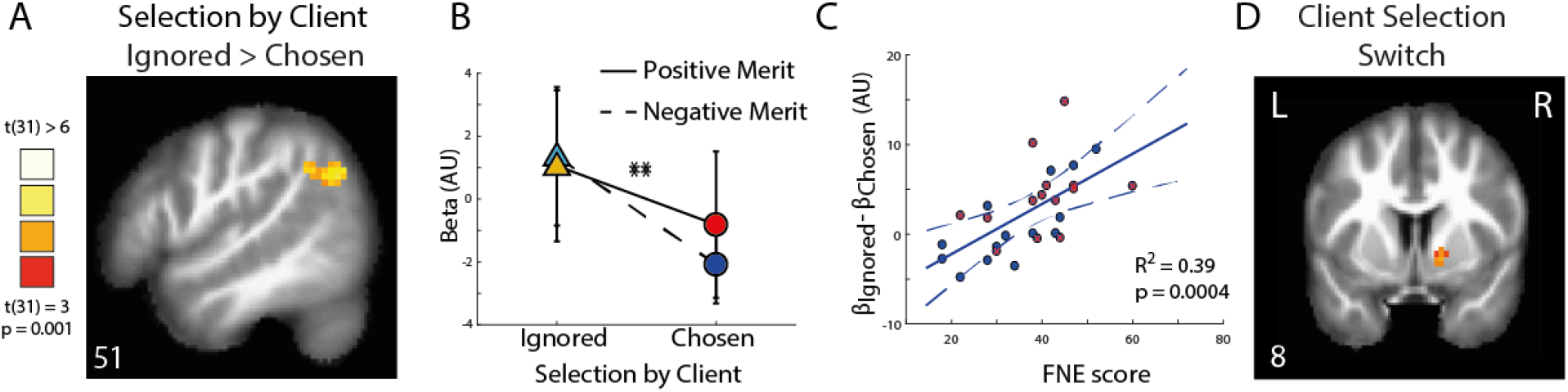
Encoding of Selection by Client and Client Selection Switch. (A) During the observation of evidence stage, activity in right TPJ (coordinates: [51,-58,35]) was higher on ignored vs chosen trials (p < 0.001, FWE cluster size corrected p < 0.05). (B) Analysis of beta estimates from the right TPJ showed a significant Selection by Client effect (N = 32, F(1,94) = 11.4, p = 0.002). Error bars indicate SEM. (C) Client selection effect on activity in the right TPJ was correlated with the participants’ fear of negative evaluation (FNE) score: differences in rTPJ activity between ignored and chosen trials was greater for participants with high FNE score (N=28, as we obtained FNE scores from a subset of the participants. Red cross indicates female participants). Regression line is computed from the population-level estimate of the FNE scores on Selection by Client effect in the rTPJ. Dashed lines indicate 2 SEs. (D) At the evidence stage activity in VS followed Client Selection Switches in adviser selection, i.e. higher when the client switched from the other adviser to you, and lower when switched from you to the other adviser (p < 0.005, FWE cluster size corrected p < 0.05, see Figure S8 for uncorrected map).

When evaluating the interaction model parameters, we observed that trait negative self-perception, as assessed by the FNE score ^8,36^, was associated with tendency to follow the defensive strategy of advice giving. Based on that observation, we hypothesised that FNE scores may predict the brain activity associated with Selection by the Client. Twenty-eight of the participants that took part in the neuroimaging experiment also completed the FNE questionnaire. Consistent with our hypothesis, we found that the rTPJ response to selection by the client (ignored > chosen, Figure 4) was modulated by participants’ individual FNE score (Figure 4C) (N=28, R^2^ = 0.39, p = 0.0004). Sensitivity of rTPJ to Selection by Client (ignored > chosen) increased with participant’s inferior self-perception, captured by the FNE scores.

To further examine the effect of Selection by Client, we used a ‘Client Selection Switch’ predictor as a regressor in a whole brain general linear model (GLM) analysis. The ‘Client Selection Switch’ predictor was set to +1 on trials when the client switched from the rival adviser to the participant, to -1 when switching from the participant to the rival adviser, and to zero when the client did not switch adviser. We found that the Ventral Striatum (VS) activity was modulated by the ‘Client Selection Switch’ variable during the evidence observation stage. VS activity increased when the client switched from the rival adviser to the participant and decreased when the client switched from the participant to the rival (Figure 4D, p < 0.001, FWE cluster size corrected p < 0.05, see Table ST3 and Figure S7 for uncorrected map). The ‘Client Selection Switch’ variable reflects the changes in the client selection, similar to the way prediction errors reflect changes in expected reward^30^, and are in line with reports of increased VS responses when selected by a client ^31^. Selection by Client was correlated with the activity in the rTPJ but changes in this variable were tracked in the VS.

At the outcome stage, participants had all the information necessary to compare the prognostic value of their own and the rival’s advice (Figure 3, Eq. [1]), and to update their Relative Merit variable by computing the Relative Merit Prediction Error (PE) (Eq. [2] and [3]). Using the model parameters evaluated for each individual, we estimated the trial-by-trial Relative Merit and Relative

Merit PE variables values, and used these as predictors in a whole brain parametric modulations analysis (see Methods). We found that during the outcome stage, activity in medial prefrontal cortex (mPFC) tracked the trial-by-trial changes in Relative Merit PE (Figure 5A, p < 0.001, FWE cluster size corrected p < 0.05, see Table ST4 and Figure S5 for uncorrected map). Given the previously documented involvement of mPFC in evaluating and inferring other agents’ traits such as reliability ^29^ and accuracy ^26,27^, and its role in mentalizing about others’ intentions and beliefs ^39^, and evaluating one’s rank in relation to others^24^, the findings reported here support the role of mPFC in social comparison with the rival adviser, tracking trial by trial the relative performance used to update relative merit.

Using the Relative Merit PE in a whole brain parametric modulation analysis during the Appraisal stage (Figure 1) we found that the VS tracked the trial by trial fluctuations in Relative Merit PE (Figure 5B, p < 0.001, FEW cluster size corrected p < 0.05, see Table ST5, and Figure S8 for uncorrected map showing bilateral VS activity). VS responses to prediction errors in social comparison are in line with previous studies showing similar VS responses to interpersonal prediction errors^40^, and to relative performance ^26^.

To further explore the temporal dynamics of tracking Relative Merit PE and Client Selection Switches, we used a region of interest (ROI) approach to examine the time courses from the mPFC and the right VS (and left VS, see Figure S8). These ROIs were defined using spheres around the coordinates indicated by NeuroSynth^41^ forward inference maps for the term ‘Ventral Striatum’, and around the peak activation in the mPFC (see Methods). We followed the steps used in previous studies^29^ to examine time courses of trials containing multiple jittered stages, aligning each trial stage timings to the stage’s mean time. We then performed a GLM on each time point across trials, in each participant, regressing the trial-by-trial Relative Merit PE and the Client Selection Switch variables from the BOLD signal and evaluating group effects (Figure 5C, D). We found that the mPFC response to Relative Merit PE, but not for Client Selection Switch, increased after the outcome stage. In the VS, responses to Relative Merit PE preceded the responses to Client Selection Switch.

**Fig. 5:**
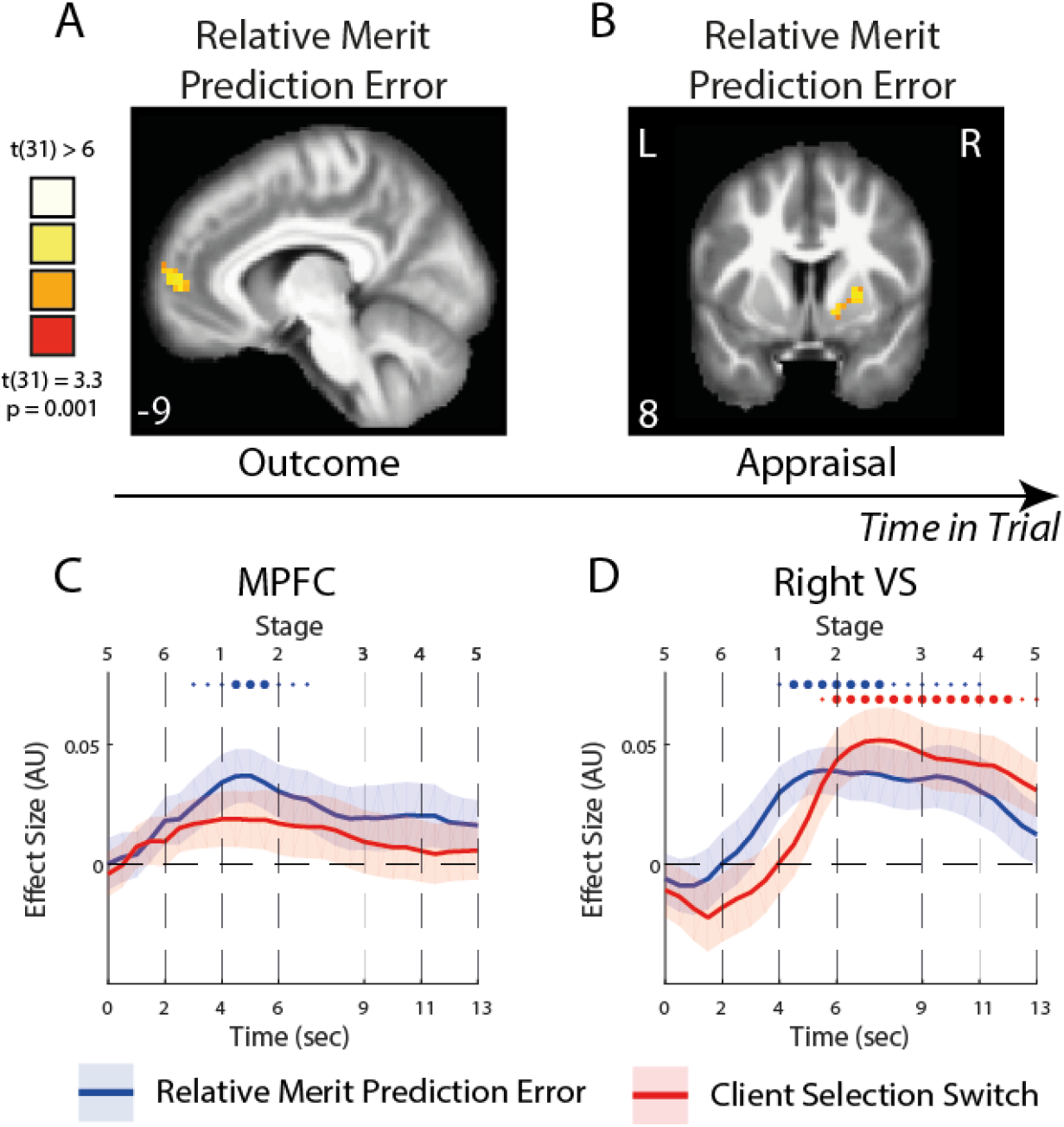
Encoding of Relative Merit Prediction Errors. (A) During the outcome stage, a whole brain analysis showed that activity in mPFC was positively modulated by the trial-by-trial Relative Merit prediction error, i.e. the differences between prognostic value comparison and previous reltive merit (Figure 2D, Eq. 1) (p < 0.001, FWE cluster size corrected p < 0.05, see Figure S7 for uncorrected map). (B) During the Appraisal stage, activity in the VS followed the Relative Merit prediction error (p < 0.001, FWE cluster size corrected p < 0.05, see Figure S8 for uncorrected map). (C) Time course of the effects of Relative Merit prediction error (blue) and Client Selection Switches (red) from the MPFC (MNI coordinates [x,y,z]: [-9,56,5]). Time courses are presented across all stages of a trial, from the showdown stage (5) of a preceding trial to the showdown stage (5) of the current trial. Thick lines indicate mean effect size, the shaddows indicate the SEM, small dots indicate p < 0.05 in these time points, and big dots indicate p < 0.005. These time courses demonstrate the temporal order of coding of Relative Merit and Selection by Client in VS. Thick lines: mean effect size. Shadows: SEM. (D) Time courses sampled from the right VS (coordinates: [18,8,-10], defined by Neurosynth), demonstrating the temporal order of coding of Relative Merit PE and Client Selection Switch.

## Discussion

We set out to study how humans attempt to influence others. We developed a novel laboratory paradigm in which participants attempted to persuade a client to take their advice over that of a competing adviser. We observed a consistent pattern of behavioural result across multiple cohorts of participants interacting with virtual or real confederates in web-based and laboratory-based experiments. Advice giving behaviour was driven by the interaction between two factors: the current level of influence over the client and the relative (i.e. social comparative) merit over the rival. Most prominently, when participants’ relative merit was positive (i.e. they were doing better than their rival), they followed what we called the ‘competitive’ strategy ^3^ by expressing higher confidence when ignored by the client and lower confidence when selected by the client. Inter-individual variability in participants’ behaviour was captured by a personalized quantitative combination of the impact of these two factors on advice confidence placing each participant’s strategy along a spectrum ranging from competitive^3^ to defensive^9^. This quantitative profile was predictive of the participant’s self-reported Fear of Negative Evaluation score^8^, thus grounding our paradigm firmly in previous research on self-esteem.

After establishing our quantitative interpretation of advice giving behaviour, we used neuroimaging to identify the neuro-computational processes proposed by our model. We found that brain activity in two distinct cortical areas consistently tracked the trial-by-trial fluctuations in ‘Relative Merit’ and ‘Selection by Client’. The mPFC tracked the Relative Merit PE, and the rTPJ tracked the fluctuations in Selection by Client. In addition, both variables were tracked in the VS. The temporal dynamics of the computations tracked by the BOLD signal in these areas were consistent with the unfolding of events across the trial, with Relative Merit PE tracked before Selection by Client.

Our study goes beyond previous investigations of the neurocognitive basis of social influence in a number of key aspects. First, it put the participants in a position to influence others’ choices. Almost all previous studies of social influence invariably concentrated on how we *react* to attempts by others to shape our behaviour (see ^20,42^ for a rare exception in which Hampton and colleagues examined strategic behaviour in the Inspector game). For example Campbell-Meiklejohnet al. ^43^ and Izuma and Adolphs^44^ investigated how our preferences change when we are given others’ opinion about objects of our desire. Zink et al.^25^ and Lignuel et al.^34^ investigated how we infer our status in a dominance hierarchy from our competitive and/or cooperative interaction with others. Although manipulation of preferences and establishing of one’s social dominance are two very common ways by which humans try to shape each other’s behaviour, these example studies did not use them as such. Instead, they focused on how the participants *reacted* to others’ attempts to shape their behaviour. For example, Ligneul et al.^34^ showed the involvement of mPFC in the dynamic representation of one’s status in a group’s hierarchy, and in the manner this information is used when choosing opponents (or avoiding them) in competitive and cooperative subsequent encounters. The alternative scenario, which was not studied, would have involved participants establishing their dominance so that trouble-making opponents would avoid them in future. To our knowledge, only one previous study has examined advice giving behaviour ^31^ directly, but again participants were not able to influence their client’s future decisions.

Second, even though many previous works studied human social interactions with more than one confederate, to our knowledge, none placed their participants in a context involving dissimilar (asymmetric) social relationships to the confederates. In many studies social interactions involved tracking another person’s reliability or intentions ^20,29^, or tracking multiple agents all playing a similar role^26–28,43^. Still other studies examined how people evaluated group behaviour, where the groups consisted of agents with similar incentives and roles and the participants engaged in various tasks such as competitive bidding ^45^, reaching consensus^46^, inferring one’s hierarchical rank ^24,34^ or tracking the group’s preferences^32^. In our task, as is the case in many real-life scenarios, the participant had to track two distinct types of relationships: a competitive one with a rival adviser, and a hierarchical one with a client whose appraisal they sought. This novel configuration of asymmetric social relationships allowed us to disentangle the separate contributions of the elements of the ‘social brain’ system ^19,22^, namely that of rTPJ and mPFC in influencing others. These contributions consisted of internally and an externally driven processes. The internal inference process was tuned to social comparison and the computation of one’s Relative Merit in the mPFC ^24,26,39^. The external process was tuned to the evaluation of social outcomes arising from others’ behaviour in relation to the self, in our case the Selection by Client, in the rTPJ. Finally, one may interpret the Selection by Client as an external event to which the participant’s attention is oriented at the beginning of every trial, in contrast, for example, to the latent process of social comparison tracked in the mPFC. The rTPJ activity elicited by external social events would therefore be in line with previous findings about the involvement of rTPJ in the orienting of attention to salient external events ^47,48^. This distinction is also in line with previous results examining strategic behaviour in two-person games ^20^, where mPFC was associated with internal inference processes and the rTPJ with evaluating external signals about the behaviour of the other.

Our findings also provide evidence for the involvement of the VS in social behaviour. Changes in both Relative Merit and Selection by Client affected ventral striatum BOLD activity, a neural structure implicated in valuation of motivational factors in decision making. This observation is in line with the combined interactive impact of these two factors on advising behaviour. Previous studies have shown that striatal neural activity may encode multiple social attributes such as reputation ^49^, selection ^25^, appraisal ^31^ and vicarious reward ^50^, and also in tracking other’s behaviour and interpersonal prediction error^40^. Here striatal activity was correlated with the different computational components of the interactive scenario as they emerged during the course of the interaction: social comparative merit and social appraisal. This finding supports the notion that the ventral striatum has a domain-general and dynamic role in valuation of various events in our environment as they occur ^30,51^.

In addition, our results indicate the importance of individual differences in social self-perception on strategic social behaviour. Computational analysis of behaviour revealed considerable variations in participants’ advising strategies, with each participant’s behaviour falling on a continuum from a pure ‘defensive’^9^ to a pure ‘competitive’^3^ strategy. These variations were captured by a personalized quantitative combination of the impact of Relative Merit and Selection by Client on advice confidence and were consistent with social rank theory: participants with negative self-perception, scoring high on fear of negative evaluation, were more likely to follow the defensive strategy^8^. In the neuroimaging experiment, participants with higher scores of negative self-perception displayed increased response in the rTPJ to being ignored by the client. These findings provide converging evidence linking management of social influence to negative self-perception. Our cognitive framework should aid future research on the social basis of mental health disorders such as depression. Deterioration of self-esteem and retreating from social engagement are two of the earliest and most debilitating hallmarks of depression^8^. The laboratory model described here offers a uniquely appropriate ecologically valid tool for measuring these social cognitive characteristics of depression.

Finally, our results demonstrate how people use confidence reports as a persuasive signal in a strategic manner. Such use of confidence reports is in line with the literature on persuasion and information sharing ^3,4,52^. However, numerous studies have used similar confidence reports to study the process of metacognition, i.e. the internal process of evaluating one’s precepts and decisions ^53,54^. Our findings demonstrate how, depending on the social context, confidence reports can depart from simply describing uncertainty about sensory information or decision variables. The findings underscore the importance of taking extra care about the framing (e.g. experimental instructions) and phrasing of *how* to ask participants to report their confidence in psychological and neuroscientific investigations. Our findings support the view of metacognition as a multi-layered process^55^, in which *expressing* one’s confidence depends not only on uncertainty about low level sensory processing or decision processes aiming to maximise reward, but also on other systematic, non-trivial sources of variance such as social comparison, closeness and friendship ^56,57^, and social expectation ^58^.

People with a more accurate opinion are often more confident. But the converse is not necessarily true: being more confident is not necessarily predictive of accuracy. Many keen observers of human condition (e.g. Bertrand Russell, W. B. Yeats and William Shakespeare, to name but a few) have complained that people who know a lot are fraught with self-doubt while the ignorant are passionately confident. Our behavioural and neurobiological results suggest that passionate overconfidence of the underdog could be better understood as a sensible recourse to the competitive strategy designed to gain higher social influence, a behaviour supported by the brain’s social and valuation systems. The philosophers’ sad lamentation therefore highlights the importance of the social comparison processes, in moderating the competitive behaviour of the ignorant.

## Acknowledgment

UH and BB are supported by the European Research Council (NeuroCoDec 309865). SP was supported by a Marie Sklodowska-Curie Individual European Fellowship (PIEF-GA-2012 Grant 328822) and is currently supported by an ATIP-Avenir grant (R16069JS) and a Collaborative Research in Computational Neuroscience ANR-NSF grant (ANR-16-NEUC-0004). CDF is supported by the Arts and Humanities Research Council (grant AH/M005933/1).

## Author Contributions

UH, BB and CDF designed the experiment. UH programmed the task. UH, SB and CO collected the data. UH and SP carried out the data analysis. UH, BB, SP and CDF wrote the manuscript.

## Methods

### Participants

We recruited four cohorts of participants for this study. All participants provided an informed consent, and received monetary compensation, both approved by the research ethics committee at UCL. Following a pilot experiment, we estimated our effect size to be around 0.5. As our experiment follows a within-participants design, we decided to recruit 60 participants for the online experiment and 30 participants for the longer lab based experiment. We recruited 60 participants for the online experiment using Amazon M-Turk. Two online participants were excluded from analysis as they did not use the full confidence scale. Online participants included 31 males (ages mean ± std 33.7 ± 9.6) and 27 females (ages 36 ± 8.5). We recruited 30 participants for a lab based experiment, in which participants carried the experiment on computers in the Psychology department building. One participant was excluded from analysis as she used only one advice level. Lab participants included 13 males (ages 26.5 ± 6) and 16 females (ages 26.2 ± 6). Finally, 34 participants were recruited for the neuroimaging part of the experiment. These corresponded to the expected effect size of activity in previous social cognition neuroimaging studies, for example ^25,31,44^. Of these, two participants were excluded from analysis due to head movements and data corruption. Neuroimaging participants therefore included 18 males (ages 24.7 ± 6.6) and 14 females (ages 23.78 ± 4.6). Finally, in the fully interactive experiment (experiment 4) we collected data from 19 triplets (57 participants, 24 males aged 25.2 ± 4.76, and 33 females aged 21.2 ± 2.25), and excluded 3 triplets from analysis. These included 2 triplets in which advisers used only the highest confidence levels (4 and 5), and one triplet in which the client chose only one adviser throughout the experiment. We therefore analysed the data from 32 advisers in the live interaction experiment.

### Client and Rival Adviser Algorithms

All participants in the main experiment played the role of an adviser, while the client and the other adviser were played by computer algorithm. The other adviser’s advice were calculated on each trial according to the probability of the coin being in the black urn (between 0-1), plus noise ((∼ *N* 0,0.08)), to range between [5W 5B], just like the participants’ advice. After outcome is revealed on each trial both advice’s prognostic value is calculated by multiplying the confidence level (1-5) by accuracy (indicating the correct coin location, i.e. W for white or B for Black urn, 1 = correct, -1 = incorrect, Eq. 1).

The client’s choice of adviser was determined by assigning an influence weight to each adviser, updating the weights after each outcome and choosing the adviser with the higher weight in the next trial. The weights summed to 10 and were set to be 5 for each adviser in the beginning of the experiment. To update the client influence weights, we used prognostic value of advice (*PA*) which were derived from advice according to the following rule: When the black urn is suggested then confidence is between [-5, -1] and *PA* is calculated by confidence +6 to range between [1, 5]. When the white urn is suggested then confidence is between [1, 5] and *PA* is calculated by confidence +5 to range between [6, 10]. We used the notation *PA*_*P*_ for to refer to prognostic value of the participant’s advice and *PA*_*O*_ for prognostic value of the other’s (rival) advice. The weights were updated after each trial according to the last trials’ prognostic values, following a rule similar to the one used by Bayarri and DeGroot ^3^:

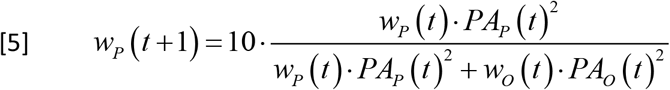

Where *w*_*P*_ is the influence weight assigned to the participant. The other adviser’s influence weight was defined as *w_O_* (*t*+1) = 10–*w_p_* (*t*+1). Note that when the influence weight of one adviser increases, the other adviser’s influence decreases in the same amount. When both advisers give the same advice the influence weights remain the same.

### Procedure

We carried out the main experiment on three different platforms: online, in the lab and in the scanner. The main experimental design features were the same across these experiments with a number of minor differences in implementation.

In the online experiment, participants were recruited using Amazon M-Turk. These participants carried out the experimental task online on their own computers using the mouse to input their confidence rating. They received a fixed monetary compensation and were promised a bonus if the client selected them on more than 100 trials. The online experiment had 130 trials. Evidence stage lasted 500ms, and all other stages of the trial were self-paced. Advice giving stage ended when confidence was reported. After the outcome was displayed, participant proceeded to the next trial by pressing a ‘Next’ button.

Lab-based participants were invited to the lab in groups of three and were told that they are about to play an adviser game together and that the roles of two advisers and client will be assigned randomly at the beginning of the experiment. The participants were then seated in isolated individual cubicles. Unbeknownst to the participants, all three players were assigned to the adviser role and played against a virtual client and a virtual rival adviser. Lab based participants received a fixed monetary compensation, and did not have any further incentive. Lab-based experiment consisted of four blocks of 70 trials. Advice giving stage was self-paced. All other stages lasted a predefined length of time: Appraisal stage lasted 1.5 seconds, evidence stage lasted 500ms, showdown stage lasted 2 seconds and outcome stage lasted 2 seconds.

In the neuroimaging experiment, participants arrived at the scanner unit and met two confederates and were given the same cover story as the lab-based participants. Participants were then put in the scanner, and were instructed on the use of response boxes for inputting their confidence ratings: left hand response box shifted the rating towards the black urn, right hand response box shifted them towards the white urn. Participants received a fixed monetary compensation. In this experiment intervals between stages were set to 1.5 second plus a jittered interval sampled from a Poisson distribution (range: 0-3 seconds; mean 1.5 second). Neuroimaging experiment consisted of four blocks with 60 trials. Advice giving stage was self-paced. All other stages lasted between 1.5 to 4.5 seconds. In addition, four intervals of 8 seconds rest were randomly dispersed between trials of each run.

In addition to the main experiment, we ran a fully interactive experiment in which all roles in the task, advisers and client, were played by human participants. This experiment was carried in a similar manner to the lab-based main experiment, with participants arriving in groups of three, and carrying the experiment on computers in separate cubicles where they were randomly assigned to the roles of advisers or client. Participants received a fixed monetary compensation and did not get any further incentive. The experiment consisted of 130 trials. The pace of the experiment depended on the interaction between the participants. Advisers waited for the client to choose one of them and then the client waited for both advisers to express their advice. See analysis of live and virtual clients’ behaviour in Supplementary Materials figure S6.

### Selective Manipulation of Advice Quality

In our early pilot sessions, we noticed that sometimes the virtual client did not shift between participants, practically ignoring one of them throughout the experiment. This happened because advisers tended to be very similarly calibrated with the evidence. As we intended to use a within-participants design to compare advice on periods in which the adviser is chosen and periods in which he is ignored by the client, we needed a way to manipulate the probability of switching between selected and ignored status. Therefore, in restricted periods of a block, we introduced some noise to one of the two advisers’ evidence such that the ratio of black and white squares in the grid became a poor predictor of the reward location. This procedure went as follows: if on a specific trial the probability of the coin being in the black urn was 0.75, the grid would normally include 75 black and 25 white squares (Figure S1). On a noisy trial with similar probability of 0.75, this composition would change to 55 black squares and 45 white squares, akin to a reduction in contrast by 20 squares. In all noisy trials contrasts were reduced by 20 squares in a similar fashion (Figure S1). The procedure ensured that one advisor’s advice accuracy was systematically inferior to the other one for a number of consecutive trials thus increasing the probability that the virtual client would shift to selecting the other adviser.

To ensure that the evidence quality manipulation was not the sole driver of changes in accuracy and difficulty throughout the task, the order of the evidence displayed to the participants in each trial (the ratio between black and white squares in the grid) was randomised for each participant. In addition, the coin location on each trial was randomly generated according to the evidence (evidence only implied the probability of the coin location). This meant that each participant experienced individual periods of high/low accuracy, which were not time locked to the evidence quality manipulation.

### Model Fitting Procedure

We used models with increased complexity to explain advice deviance reported by the participants. The most elaborated model is the *interaction model*, described in equation [4], which assumes that advice deviance is affected by: systematic bias in confidence (which is in fact the intercept or alpha parameter), current Selection by Client (ignored/chosen), Relative Merit tracked by comparison with rival (Eq. [3]), and the interaction between Relative Merit and Selection by Client. Simpler models included: a *mixture model* excluding the interaction parameter, a Relative Merit *model* excluding all selection parameters, a *selection model* excluding all Relative Merit parameters, and a *bias model* excluding all Relative Merit and Selection by Client parameters. An additional model was also tested that was identical to the ‘Interaction’ model but used the magnitude and sign of Relative Merit instead of only the sign of Relative Merit.

We fitted all models to individual advice deviance. We used a cost function, *L(M)* to estimate a given model M fit to the data. The cost function compared the advice deviance estimated with the model M (*AdviceDeviance_M_* (*t*)) and the actual advice deviance observed in behaviour on each trial (*AdviceDeviance_Data_ (t))*:

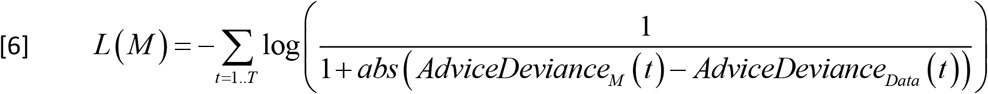

Like log likelihood cost function, the ratio inside the log is close to one when the estimation is close to the data, and it gets closer to zero when the distance between estimation and data increases. Therefore, lower values of the cost function indicate better fit of the model to the data. We used a Markov Chain Monte Carlo (MCMC) Metropolis-Hastings algorithm for model fitting and estimation for each participant ^59–61^. For model comparisons we calculated individual Deviance Information Criterion (DIC) ^60^, which uses the distribution of likelihood obtained and penalizes for increased number of parameters (Supplementary Figure S2). We used in house Matlab code and the MCMC toolbox for Matlab by Marko Laine (http://helios.fmi.fi/∼lainema/mcmc/#sec-4).

Parameter estimation was done individually by integrating the marginal distribution of the parameter values, uncovered using the Markov process chain ^61^. The learning rate parameters estimated using the Interaction model were used in the aggregated analysis to determine the trial by trial Relative Merit and separate trials to positive and negative Relative Merit. The mean parameter estimations for all models are reported in table ST1 in the supplementary materials.

### MRI data acquisition

Structural and functional MRI data were acquired using Siemens Avanto 1.5 T scanner equipped with a 32-channel head coil at the Birkbeck-UCL Centre for Neuroimaging. The echoplanar image (EPI) sequence was acquired in an ascending manner, at an oblique angle (≈30°) to the AC–PC line to decrease the impact of susceptibility artefact in the orbitofrontal cortex ^62^ with the following acquisition parameters: volumes, 44 2mm slices, 1mm slice gap; echo time = 50 ms; repetition time = 3740 ms; flip angle = 90°; field of view = 192 mm; matrix size = 64 × 64. As the time for each block was dependent on the participants’ reaction time, overall functional blocks changed in length, and approximately 250 volumes were acquired in about 15 minutes and 40 seconds. A structural image was collected for each participant using MP-RAGE (TR = 2730 ms, TE = 3.57 ms, voxel size = 1 mm3, 176 slices). In addition, a gradient field mapping was acquired for each participant.

### fMRI data analysis

Imaging data were analysed using Matlab (R2013b) and Statistical Parametric Mapping software (SPM12; Wellcome Trust Centre for Neuroimaging, London, UK). Images were corrected for field inhomogeneity and corrected for head motion. They were subsequently realigned, coregistered, normalized to the Montreal Neurological Institute template, spatially smoothed (8 mm FWHM Gaussian kernel), and high filtered (128 seconds) following SPM12 standard preprocessing procedures.

We carried two complementary data analyses. Using the computational behaviour analysis, we labelled trials according to Selection by Client (chosen/ignored) and Relative Merit (positive/negative). We examined the effect of Selection by Client, Relative Merit and their interaction on brain activity. We used individual level general linear model (GLM) with stick predictors at the onset of each stage (appraisal, evidence, advice report, other advice display, outcome), and additional boxcar predictor 1.5 seconds long ending at the time of advice report confirmation, capturing the motor button presses. We ran GLMs in which the condition labels (Selection by Client (chosen/ignored) and Relative Merit (positive/negative)) applied to the outcome stage, appraisal stage and evidence stage separately, to overcome the problem of correlation between predictors of interest, as the order of our stages was fixed. In addition we ran a whole brain parametric modulation analysis with a ‘Client Selection Switch’ predictor, set to 1 when the client switched from the rival adviser to the participant, -1 when switching from the participant to the rival adviser, and 0 when the client made the same choice as before.

In a separate analysis we examined how the trial-by-trial Relative Merit PE modulated brain activity. We used a set of variables as parametric modulators of activity which included the Relative Merit PE as the regressor of interest, and aggregated Relative Merit, the trial-by-trial prognostic value comparison, client’s reward prediction error, and unsigned participant’s advice as regressors of no interest. We used different GLMs to estimate the effect of this set of parameter modulations on the outcome stage, appraisal stage and evidence stage separately.

In region of interest (ROI) analysis we examined event related effects in specific brain region in different stages within an advice giving trial. Individual beta maps were estimated and sampled within ROIs using MarsBar SPM toolbox. We used NeuroSynth^41^ defined ROIs of the left and right ventral striatum. We selected the peak of the ventral striatum reverse inference map, made from 310 studies. We used a 12mm sphere around the left and right peak activity as ROI using MarsBar SPM toolbox (MNI coordinates [-12 8 -8], [10 6 -8], z > 22).

To examine the time course of the changes in brain activity in the regions of interest, we followed previous studies^29^ and exploited the time jitters to disentangle the brain activity corresponding to different cognitive processes of interest. We separated each subject’s time series sampled from the VS into each trial, and resampled each trial to a duration of 15s, aligned according to the trial stages (Figure 1): previous trial’s showdown(stage 5) at time 0, previous trial’s outcome (stage 6) at time 2s, appraisal (stage 1) at time 4s, evidence (stage 2) at time 6s, confidence report (stages 3-4) at time 9s-11s, and current trial’s showdown (stage 5) at time 13s (These timings were the mean timings across all trials in all subjects.) The resampling resolution was 100ms. This temporal realignment allowed the observation of signal throughout the trial while taking advantage of the random jitter and fast event-related design. We then performed a GLM at each time point across trials in each subject. We had one regressor for ‘Relative Merit PE’, and another for ‘Client Selection Switch’. We then calculated the mean of the effect across subjects at each time point, and their standard errors.

### Questionnaires

As follow up for our experiment we sent participants Fear of Negative Evaluation (FNE) questionnaire ^36^ to be filled online, to test the link between trait selection perception and social behaviour, as predicted by social rank theory ^35^. The questionnaire was sent six months after the main experiment, to make sure there is no effect of the questionnaires on the performance in the task and vice versa. Participants were paid for completing the questionnaires. 69 of our original 120 participants filled the questionnaire, 29 from the fMRI cohort, 15 from the lab cohort and 25 from the online cohort.

### Data Availability

The behavioural data that support the findings of this study are available from figshare (https://doi.org/10.6084/m9.figshare.5414350.v1) ^63^. The statistical parametric maps from the neuroimaging part of the experiment are available from NeuroVault: http://neurovault.org/collections/2204/.

